# Kidins220 sets the threshold for survival of neural stem cells and progenitors to sustain adult neurogenesis

**DOI:** 10.1101/2023.01.10.523252

**Authors:** Ana del Puerto, Beatriz Martí-Prado, Ana L. Barrios-Muñoz, Coral López-Fonseca, Julia Pose-Utrilla, Berta Alcover-Sanchez, Fabrizia Cesca, Giampietro Schiavo, Miguel R Campanero, Isabel Fariñas, Teresa Iglesias, Eva Porlan

## Abstract

In the adult mammalian brain, neural stem cells (NSCs) located in highly restricted niches sustain the generation of new neurons that integrate into existing circuits. A reduction in adult neurogenesis is linked to ageing and neurodegeneration, whereas dysregulation of proliferation and survival of NSCs have been hypothesized to be at the origin of glioma. Thus, unravelling the molecular underpinnings of the regulated activation that NSCs must undergo to proliferate and generate new progeny is of considerable relevance. current research has identified cues promoting or restraining NSCs activation. Yet, whether NSCs depend on external signals to survive or if intrinsic factors establish a threshold for sustaining their viability remains elusive, even if this knowledge could involve potential for devising novel therapeutic strategies. Kidins220 (Kinase D-interacting substrate of 220 kDa) is an essential effector of crucial pathways for neuronal survival and differentiation. It is dramatically altered in cancer and in neurological and neurodegenerative disorders, emerging as a regulatory molecule with important functions in human disease. Herein, we discover severe neurogenic deficits and hippocampal-based spatial memory defects in Kidins220 deficient mice. Mechanistically, we demonstrate that Kidins220-dependent activation of AKT in response to EGF restraints GSK3 activity preventing NSCs apoptosis. Hence, Kidins220 levels set a molecular threshold for survival in response to mitogens, allowing adult NSCs to proliferate. Our study identifies Kidins220 as a key player for sensing the availability of growth factors to sustain adult neurogenesis, uncovering a molecular link that may help paving the way towards neurorepair.

## INTRODUCTION

In the mammalian adult brain, two specialized microenvironments contribute to the maintenance of neural stem cell (NSC) pools to sustain continuous neurogenesis throughout life. In these niches, NSCs and their own cellular progeny coexist with supporting cells, a specialized extracellular matrix and vasculature. The mouse subependymal zone (SEZ) comprises a layer of cells juxtaposed to the ventricular wall lined by ependymal cells, amidst which residing NSCs extend a primary cilium to contact the cerebrospinal fluid and sense the milieu. In the SEZ, relatively quiescent NSCs (B cells) give rise to fast-dividing intermediate progenitor cells (C cells). Arising from these progenitors are neuroblasts (type A cells) that migrate into the olfactory bulb (OB), to differentiate into several types of interneurons, that contribute to refine the processing of olfactory information [1]. Similarly, in the subgranular zone (SGZ) of the hippocampal dentate gyrus (DG), NSCs termed radial glia-like (RGL)-type 1 cells [2] give rise to intermediate progenitors (type 2a and 2b cells), which generate new neurons that migrate into the DG cell layer where they integrate into hippocampal circuits [3].

The generation of new neurons as an ongoing dynamic process echoes in adulthood the mechanisms driving neurogenesis during neurodevelopment. The number of NSCs and newly generated neurons can be modulated at all phases of the process by restraining or promoting proliferation, differentiation, migration, survival, maturation, and integration of newborn neurons into the existing circuitry [3, 4].

Stress, aging, and neurodegeneration are notable negative regulators of adult neurogenesis involved in memory loss, mood alterations and additional behavioral changes, as revealed by studies in rodents (reviewed in [5]). In contrast, dysregulation of proliferation and survival of NSCs have been hypothesized to be at the origin of glioma (reviewed in [6, 7]). Thus, dissecting the molecular basis of the regulated transition that NSCs must undergo from quiescence to activation to generate new progeny is of considerable relevance since it bestows potential for devising novel stem cell-based therapeutic strategies. At the molecular level, in NSCs this transition is governed by numerous factors, either inherent to them or to the niche [3, 4], and even systemic to the entire organism [8]. Although short and long-distance cues have shown effects in the activation of adult NSCs, little is known about how their survival, serving as a requisite for proliferation, is regulated. Whether NSCs are dependent on external signals to survive or whether intrinsic factors establish a threshold for sustaining the viability of these cell populations is unclear [9].

Kidins220 (Kinase D-interacting substrate of 220 kDa) [10], also known as ARMS (Ankyrin repeat-rich membrane spanning) [11], is an essential gene whose roles keep emerging. It functions as a signaling scaffold at the plasma membrane as an effector of several signaling pathways. Kidins220 acts downstream of neurotrophin receptors and interacts with diverse signaling pathways to promote neuronal survival, differentiation, and synaptic activity [12, 13], and acts as a regulator of nervous system development [14, 15]. KIDINS220 expression is altered in cancer and in neurological and neurodegenerative disorders, including cerebral ischemia, Alzheimer’s disease, Huntington’
ss disease and idiopathic normal pressure hydrocephalus [16-23]. Additionally, rare novel missense and loss-of-function variants in *KIDINS220* gene are associated with schizophrenia [24, 25] and, more recently, SINO (spastic paraplegia, intellectual disability, nystagmus, and obesity) syndrome and ventriculomegaly [26-28], altogether identifying KIDINS220 as a multifaceted player with important functions in human disease.

Herein, we identify Kidins220 as a novel intrinsic regulator of NSCs in adult neurogenic niches, and report that its decreased expression provokes severe neurogenic deficits and hippocampal-based spatial memory defects. Mechanistically, we demonstrate that Kidins220-dependent activation of AKT in response to epidermal growth factor (EGF) restraints glycogen synthase kinase-3 (GSK3) activity preventing NSCs apoptosis. Our study identifies Kidins220 as a key player for sensing the availability of growth factors to sustain adult neurogenesis, uncovering a molecular mechanism by which Kidins220 confers NSCs responsiveness to mitogens, setting a molecular threshold for NSC survival.

## RESULTS

### Kidins220 deletion in adult neural stem cells from the subependymal zone

To examine Kidins220 expression by NSCs in the SEZ, we combined antibodies against Kidins220 [10, 20, 23], sex determining region Y-box 2 (Sox2) transcription factor (expressed by B1 cells and by progenitors [29] and glial fibrillar acidic protein (GFAP; expressed by B1 stem cells [3]. In the SEZ, Kidins220 immunostaining was particularly bright in GFAP^+^-Sox2^+^ astrocytes localized in the vicinity or in direct contact with the walls of the lateral ventricle (Fig. 1A). SEZ GFAP^+^ NSCs span amongst multiciliated ependymal cells, exposing a small portion of their membrane with a primary cilium to the ventricle lumen, in so-called ‘pinwheel’ structures. Ciliary (both motile and primary) basal bodies can be labelled with antibodies against γ-tubulin [30]. As shown (Fig. 1B), anti-Kidins220 antibodies stained very strongly NSCs, i.e., GFAP^+^-cells with a primary cilium (γ-tubulin^+^), located within a rosette of multiciliated ependymal cells, labeled with N-Cadherin [30, 31]. *Kidins220*-floxed (*Kidins220*^*f/f*^) mice have been described previously as a hypomorphic model of Kidins220 and display reduced levels of Kidins220 when compared to control mice in different tissues [23]. SEZ homogenates from *Kidins220*^*f/f*^ also showed Kidins220 protein downregulation (Fig. 1C-E). To investigate whether low Kidins220 levels affect adult NSCs and neurogenesis, we injected 2 months-old wild-type and *Kidins220*^*f/f*^ mice with the thymidine analogue 5-chloro-2’-deoxyuridine (CldU) and kept them alive for 4 weeks, to label slow dividing NSCs at the SEZ, and new neurons at the OB, by CldU-retention. Constitutively reduced Kidins220 levels during development did not provoke adult NSC decreased seeding, since total number of NSCs, scored as CldU^+^-label retaining cells in the SEZ were unaltered in *Kidins220*^*f/f*^ *vs*. wild-type mice (Fig. 1F, G). To obtain a complete deletion of Kidins220 in adult NSCs, we crossed *Kidins220*^*f/f*^ animals with mice expressing the Cre recombinase under the mouse *Gfap* promoter [32], to produce *Kidins220*^*Gfap*Δ/Δ^ mice. Immunoblot in cultured primary astrocytes from wild-type, *Kidins220*^*f/f*^ and *Kidins220*^*GfapΔ/Δ*^ mice showed complete Kidins220 ablation (Fig. S1), and Kidins220 was undetectable in GFAP^+^ cells from *Kidins220*^*GfapΔ/Δ*^ mice in the SEZ (Fig. 1E). As for *Kidins220*^*f/f*^, we did not find differences in the total number of NSCs in the SEZ of *Kidins220*^*GfapΔ/Δ*^ mice (Fig. 1F). Accordingly, no changes were detected in NSCs/progenitors labelled with Sox2, in the neuronal-lineage marker doublecortin (DCX)^+^-neuroblast population, nor in the newly generated OB neurons (scored as CldU^+^ cells at the OB glomeruli) (Fig. S2A-C). Likewise, newly incorporated oligodendrocytes to the corpus callosum (CldU^+^ cells), and newly formed terminally differentiated astrocytes at the SEZ (CldU^+^-S100β^+^-double positive cells) [33] were unaltered (Fig. S2D-E), indicating that Kidins220 deficiency does not alter NSC-astrocytic terminal differentiation nor oligodendrogenesis. Kidins220 has been recently involved in the etiopathology of hydrocephalus, as *Kidins220*^*f/f*^ mice present ventriculomegaly at different degrees [23]. Similarly to floxed mice, *Kidins220*^*Gfap*Δ/Δ^ mice presented enlarged ventricles, as measured by magnetic resonance imaging (MRI), when compared to wild-type littermates (Fig. S3). To directly examine NSCs at the ventricular surface of Kidins220 mutants, we used whole mount *en face* staining of NSCs. As previously described for *Kidins220*^*f/f*^ [23], *Kidins220*^*Gfap*Δ/Δ^ ventricular wall linings were apparently normal, showing multiciliated ependymal cells (Fig. 1G). In addition, we were able to detect NSCs exposed at the ventricle in *Kidins220*^*f/f*^ and *Kidins220*^*Gfap*Δ/Δ^ mice, with no overall changes in the pinwheel organization (Fig. 1G).

**Figure 1.**
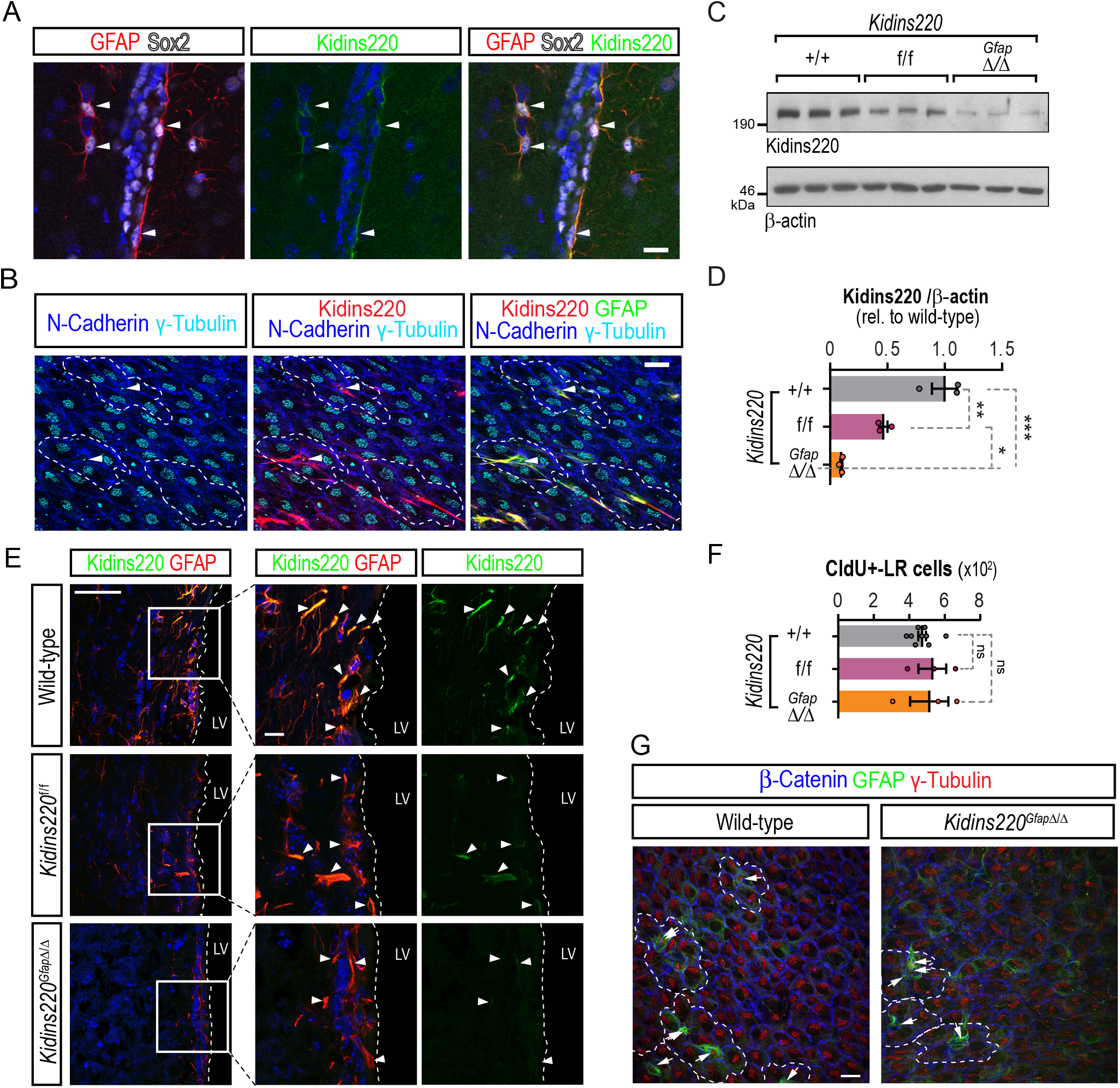
Kidins220 deletion in adult neural stem cells from the subependymal zone. **A**, Confocal micrographs of the staining for GFAP (red), Sox2 (gray) and Kidins220 (green) in the subependymal zone (SEZ) of wild-type mice. Triple-positive cells are shown (white arrowheads). Nuclei are stained with DAPI (blue). Scale bar 20 μm. **B**, Representative confocal micrographs of lateral ventricle wholemount *en face* preparations stained for N-cadherin (blue), γ-tubulin (cyan), Kidins220 (red) and GFAP (green) to reveal the characteristic pinwheel structure (white dashed lines) where SEZ stem cells reside (white arrowheads). Scale bar 20 μm. **C**, Representative Kidins220 and ß-actin (loading control) immunoblots of SEZ lysates from wild-type, *Kidins220*^*f/f*^ and *Kidins220*^*GfapΔ/Δ*^ mice. Each lane represents extracts from one mouse. **D**, Kidins220 levels represented in arbitrary units after normalization with ß-actin in wild-type, *Kidins220*^*f/f*^ and *Kidins220*^*GfapΔ/Δ*^ mice. Data represent mean ± s.e.m.; each data point represents an individual mouse (N=3, for each condition). *P< 0.05, **P < 0.01, ***P < 0.001, by one-way ANOVA followed by Tukey’s *post-hoc* test. **E**, Confocal micrographs of the staining for GFAP (red) and Kidins220 (green) in the SEZ of wild-type, *Kidins220*^*f/f*^ and *Kidins220*^*GfapΔ/Δ*^ mice. Nuclei are stained with DAPI (blue). Scale, 50 μm (left), 10 μm (insets). LV, lateral ventricle (delimited by white dashed lines). White arrowheads point at GFAP positive cells. **F**, Quantification of the numbers of CldU^+^-label retaining (LR) cells at the lateral ventricles of wild-type (N=9), *Kidins220*^*f/f*^ (N=3) and *Kidins220*^*Gfap*Δ/Δ^ (N=3) mice. **G**, Representative confocal images of wild-type and *Kidins220*^*Gfap*Δ/Δ^ lateral ventricle wholemount preparations stained for ß-catenin (blue), γ-tubulin (red) and GFAP (green) to reveal pinwheels (white dashed lines) and SEZ stem cells (arrows). Scale bar: 10 μm.

### Neurogenic defect in the hippocampus of Kidins220 deficient mice

We next inspected the SGZ. Immunodetection of Sox2 and GFAP to label RGL-type 1 cells combined with antibodies against Kidins220 showed that it was abundant in the SGZ, labelling NSCs (Fig. 2A, B). We found decreased expression in GFAP^+^ cells in *Kidins220*^*f/f*^ tissue, and below detection threshold in GFAP^+^ cells from *Kidins220*^*GfapΔ/Δ*^ mice in the SGZ (Fig. 2C), which was confirmed in hippocampal tissue extracts by immunoblot (Fig. 2D, E). Histological examination of *Kidins220*^*f/f*^ and *Kidins220*^*GfapΔ/Δ*^ DG showed that cytoarchitectural organization was apparently preserved, although the volume was significantly smaller than that of wild-type mice (Fig. 2F, G). Similarly to the SEZ, Kidins220 deficiency did not alter the total number of NSC, scored as Sox2^+^-CldU^+^ cells, (Fig. 2H). However, we discovered a sharp decrease in the numbers of total CldU^+^ cells in the DG (Fig. 2I), suggesting a deficit in newborn neurons. By combining Sox2 and 5-iodo-2’-deoxyuridine (IdU) stainings (injected one hour before sacrifice to label proliferative progenitors [31]), we found a decrease in type 2 cells in Kidins220-deficient mice (Fig. 2J, K). This reduction was not due to impaired proliferation of progenitors, since the rate of cycling (Ki67^+^)-Sox2^+^ cells was unaffected in *Kidins220*^*f/f*^ and *Kidins220*^*GfapΔ/Δ*^ compared to wild-type mice (Fig. 2L). Concomitantly, we found a reduction in the number of neuroblasts expressing Dcx in *Kidins220*^*Gfap*Δ/Δ^ mice (Fig. 2M, N), albeit this decrease did not reach significance in *Kidins220*^*f/f*^ animals (Fig. 2M, N). We then determined the emergence of newborn neurons in mutant SGZs, scoring the percentage of CldU^+^ cells that had acquired the neuronal marker NeuN. We observed a sharp decrease in newly generated neurons in *Kidins220*^*f/f*^ and *Kidins220*^*GfapΔ/Δ*^ mice, indicating a neurogenic deficit in adulthood (Fig. 2O, P).

**Figure 2.**
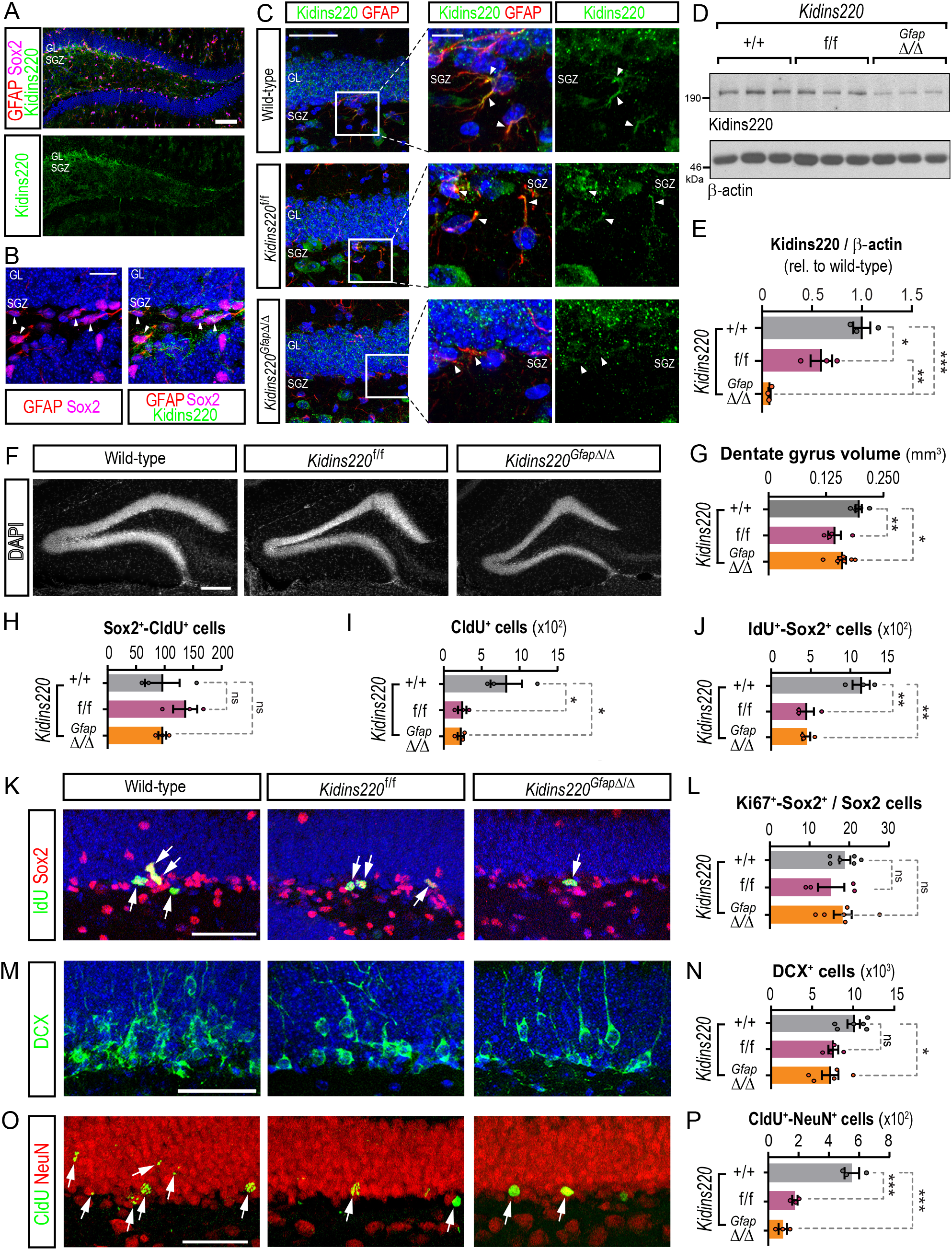
Neurogenic defect in the hippocampus of Kidins220 deficient mice. **A**, Confocal micrographs of the staining for GFAP (red), Sox2 (magenta) and Kidins220 (green) in the dentate gyrus (DG) of wild-type mice. Nuclei are stained with DAPI (blue). Scale bar 100 μm. **B**, Higher magnification confocal micrograph of the staining for GFAP (red), Sox2 (magenta) and Kidins220 (green) showing specific co-staining in cells of the subgranular zone (SGZ) of wild-type mice (white arrowheads). Nuclei are stained with DAPI (blue). Scale bar 20 μm. **C**, Confocal micrographs of the staining for GFAP (red) and Kidins220 (green) in the DG of wild-type, *Kidins220*^*f/f*^ and *Kidins220*^*GfapΔ/Δ*^ mice. Nuclei are stained with DAPI (blue). Scale, 50 μm (left), 10 μm (insets). GL, granular zone; White arrowheads point at GFAP^+^ cells. **D**, Representative Kidins220 and ß-actin (loading control) immunoblots of hippocampal lysates from wild-type, *Kidins220*^*f/f*^ and *Kidins220*^*GfapΔ/Δ*^ mice. Each lane represents extracts from one mouse. **E**, Kidins220 levels represented in arbitrary units after normalization with ß-actin in hippocampal lysates in wild-type, *Kidins220*^*f/f*^ and *Kidins220*^*GfapΔ/Δ*^ mice (N=3, each). Data represent mean ± s.e.m.; each data point represents an individual mouse. *P< 0.05, **P < 0.01, ***P < 0.001, by one-way ANOVA followed by Tukey’s *post-hoc* test. **F**, Confocal micrographs of the DG of wild-type, *Kidins220*^*f/f*^ and *Kidins220*^*Gfap*Δ/Δ^ mice. Nuclei are stained with DAPI (gray). Scale bar: 200 μm. **G**, Quantification of the DG volume (mm^3^) in wild-type (N=5), *Kidins220*^*f/f*^ (N=4) and *Kidins220*^*Gfap*Δ/Δ^ (N=7) mice. **H**, Numbers of Sox2^+^-CldU^+^ cells, **I**, CldU^+^ cells and **J**, IdU^+^-Sox2^+^ cells in the SGZ of wild-type, *Kidins220*^*f/f*^ and *Kidins220*^*Gfap*Δ/Δ^ mice (N=3, for each condition). **K**, Representative confocal images of the co-staining for Sox2 (red) and IdU (green, arrows). Nuclei are stained with DAPI (blue). Scale bar: 50 μm. **L**, Percentage of Sox2^+^ cells positive for Ki67 in the SGZ of wild-type (N=6), *Kidins220*^*f/f*^ (N=4) and *Kidins220*^*Gfap*Δ/Δ^ (N=6) mice. **M**, Confocal images of the staining for the neuroblast marker doublecortin (DCX, green) and DAPI (blue). Scale bar: 50 μm. **N**, Quantification of the total numbers of DCX^+^ cells in wild-type (N=6), *Kidins220*^*f/f*^ (N=4) and *Kidins220*^*Gfap*Δ/Δ^ (N=6) mice. **O**, Representative confocal images of the staining for the neuronal marker NeuN (red) and CldU (green) 3 weeks after injection to label new-born neurons. Scale bar: 50 μm. **P**, Quantification of CldU^+^-NeuN^+^ double positive cells in wild-type, *Kidins220*^*f/f*^ and *Kidins220*^*Gfap*Δ/Δ^ mice (N=3, each). Data represent mean ± s.e.m.; each data point represents an individual mouse. ns, not significant, \**P* < 0.05, \*\**P* < 0.01, \*\*\**P* < 0.001, by one-way ANOVA and Dunnett’s *post-hoc* test.

### Functional consequences of Kidins220 deficiency in hippocampal neurogenesis

To analyze the functional consequences of the reduced adult hippocampal neurogenesis, we performed behavioral tests. To assess hippocampal-based spatial working memory, mice from the 3 genotypes were subjected to the T maze test (Fig. 3A, B). For the parameters scored during the first week (latency, accuracy, and visits to the unrewarded arm), results showed there was not a statistically significant interaction between the effects of genotype and sessions (Fig. 3A and see Table S3 for detailed ANOVA results). As expected, the variable ‘genotype’ did not have a statistically significant effect on spatial working memory whilst, the variable ‘session’ did, for all three parameters scored. This indicates that during acquisition phase, all mice independently of their genotypes, showed a similar reduction in the number of wrong visits and latency, and a similar increase in accuracy, indicating a successful acquisition of T-maze discrimination learning throughout the sessions (Fig. 3A). In contrast, ‘genotype’ and ‘session’ had a statistically significant impact on reversal learning for the three parameters scored (Fig. 3A). This indicated that during the reversal phase, only wild-type mice showed a successful reversal learning, obtaining a good score of accuracy and a reduction of wrong visits and latency compared to *Kidins220*^*f/f*^ and *Kidins220*^*GfapΔ/Δ*^ mice. A closer examination of the data suggested that wild-type mice successfully achieved learning on the third session. In contrast, both *Kidins220*^*f/f*^ and *Kidins220*^*GfapΔ/Δ*^ mice displayed an increased latency in completing the trial and in the number of visits to the wrong arm, while showed a lack of accuracy in comparison with wild-type mice in session 3 (Fig. 3B). Taken together, these results suggest that physiological Kidins220 levels promote cognitive flexibility.

**Figure 3.**
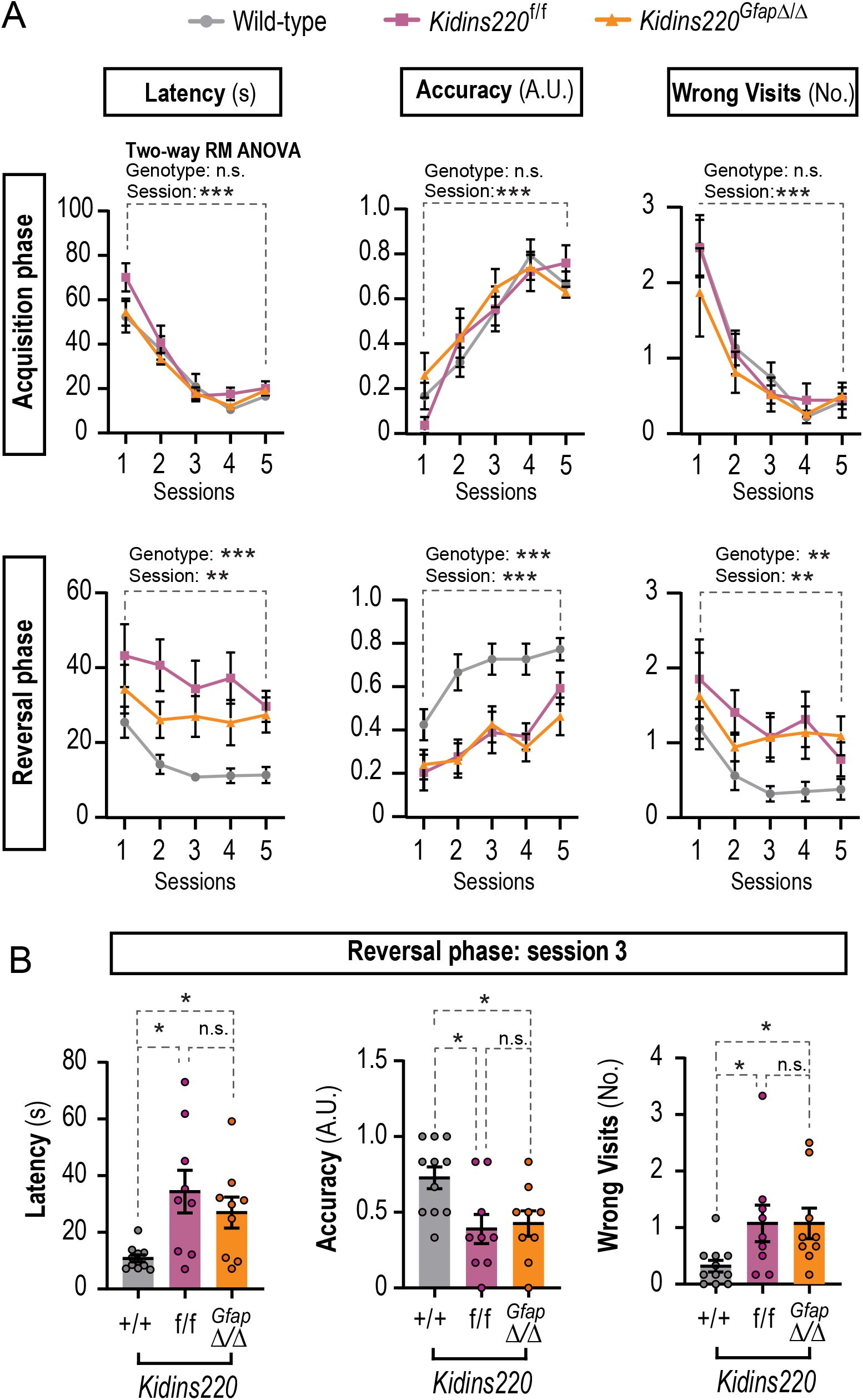
Kidins220-defficient mice exhibit severe deficits in spatial memory. **A**, T maze test results during acquisition phase (upper panels) and reversal phase (lower panels). Wild-type (N=11), *Kidins220*^*f*/f^ (N=9) and *Kidins220*^*GfapΔ/Δ*^ (N=9) mice were challenged for 5 sessions, with 6 trials in each session, and three parameters were scored: latency in rewarded arm (the time (s) to reach the reward and start eating), accuracy (arbitrary units: 1 for entering the rewarded arm and eating from the pellet within 30 seconds; 0 for any other outcome), and the number of visits to the non-rewarded arm (wrong visits). The effects of the genotype and sessions and their interaction were analyzed by two-way repeated measures (RM) ANOVA. Interaction was non-significant (*P* > 0.05). **B**, One-way ANOVA analyses of data from session 3 in the reversal phase, for latency, accuracy and number of wrong visits for wild-type, *Kidins220*^*f*/f^ and *Kidins220*^*GfapΔ/Δ*^ mice. Data represent mean ± s.e.m.; each data point represents an individual mouse. Ns, not significant, \**P* < 0.05, \*\**P* < 0.01, *** *P* < 0.001, by two-way RM (A) or one-way ANOVA followed by Dunnett’s T3 (latency) or Tukey’s *post-hoc* tests (accuracy and wrong visits).

### Kidins220 deficiency in NSC decreases cell survival

Fresh tissue from both neurogenic regions can be dissected, homogenized and cultured *in vitro* as free-floating aggregates known as neurospheres [34, 35], or alternatively grown in adherent conditions seeded on a laminin-rich substrate [36]. In both cases, in the presence of epidermal growth factor (EGF) and fibroblast growth factor-2 (FGF-2) as main mitogens, NSCs derived from the SEZ and the DG thrive and proliferate over several passages maintaining their multipotency. Importantly, NSCs cultured *in vitro* expressed detectable levels of Kidins220 (Fig. S4A). Furthermore, hippocampal NSCs obtained from *Kidins220*^*f/f*^ and *Kidins220*^*GfapΔ/Δ*^ grown *in vitro* yielded significantly less primary neurospheres than wild-type cells (Fig. 4A), consistent with the observed defect in NSCs and progenitors in the SGZ. Accordingly, attempts of expanding these cultures *in vitro* failed, corroborating *in vivo* data showing the extreme sensitivity of the SGZ to Kidins220 levels. As performing *in vitro* experiments using NSCs from the DG of Kidins220 deficient mice proved impossible, we established NSC cultures from the SEZ. *Kidins220*^*f/f*^ neurospheres expressed half the levels than wild-type, while Kidins220 levels were undetectable in *Kidins220*^*GfapΔ/Δ*^ cells (Fig. S4B). Despite the detection of normal numbers of NSCs in the intact SEZ of *Kidins220*^*GfapΔ/Δ*^ mice, we obtained reduced numbers of neurosphere-initiating cells (Fig. 4B) indicating a cell autonomous defect, not obvious *in vivo* at the age studied. In contrast, *Kidins220*^*f*/*f*^ SEZs did not display obvious differences in primary neurosphere output (Fig. 4B), suggesting that a certain level of Kidins220 is required for NSCs to thrive *in vitro* (Fig. S4B). *Kidins220*^*GfapΔ/Δ*^ SEZ cells could be passaged but yielded reduced accumulated cell numbers *vs*. wild-type and *Kidins220*^*f/f*^ cells (Fig. 4C). This phenotype was accompanied by a marked reduction of neurosphere diameter (Fig. 4D, E) with no substantial defect in proliferation rate (Fig. 4E, F), altogether suggesting a viability defect. Staining for 7-aminoactinomycin D (7AAD) and Annexin V, to assess the percentage of live and apoptotic cells in NSC cultures by flow cytometry, showed that *Kidins220*^*Gfap*Δ/Δ^ cells had an overall decreased survival and a higher rate of apoptosis than wild-type, affecting as much as the 65% of the culture, hence suggesting a cause for NSC depletion. (Fig. 4G-I).

**Figure 4.**
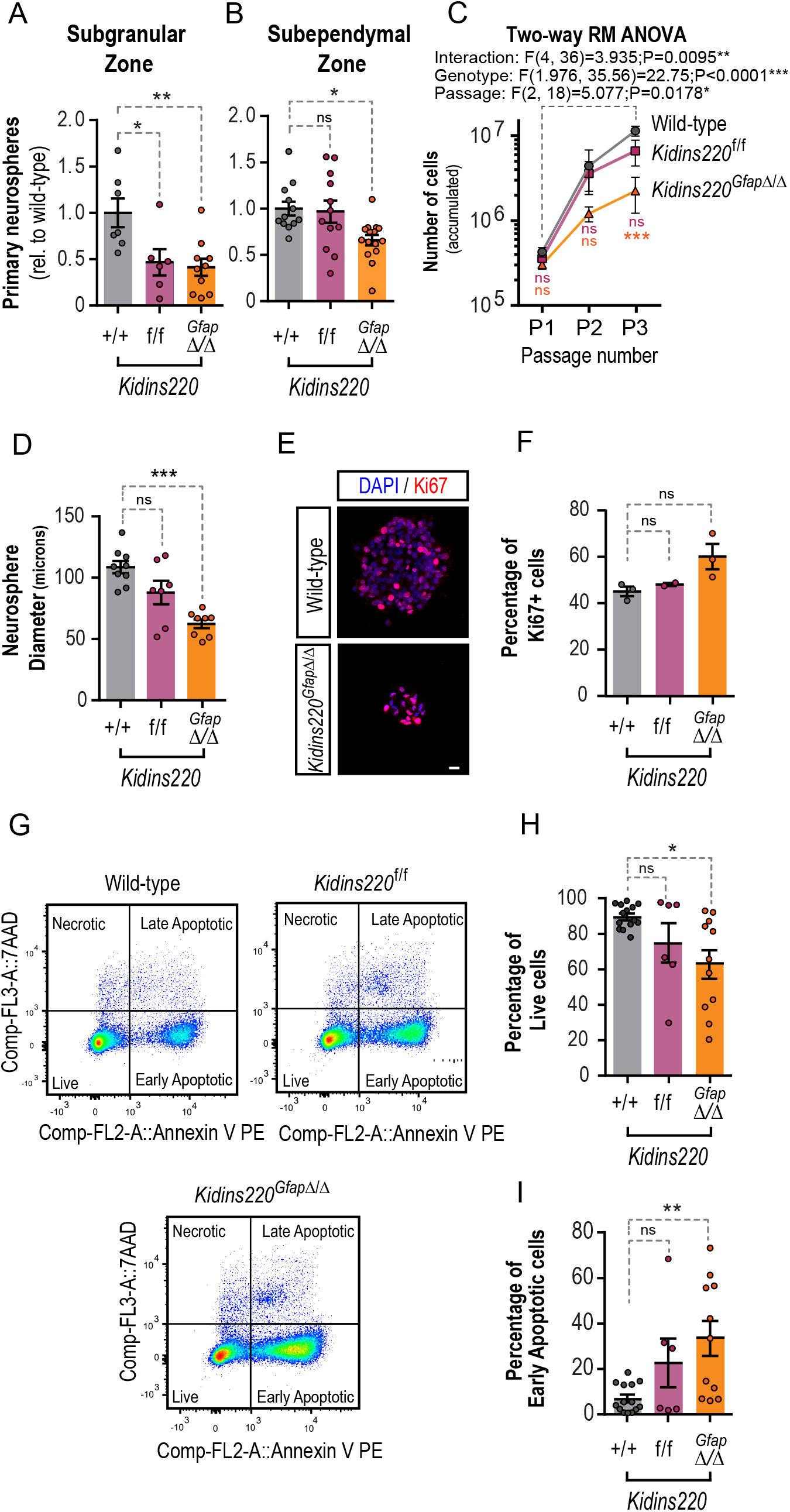
Viability defect in Kidins220-deficient neural stem cells. Primary neurospheres obtained from the hippocampus (**A**) and the walls of the lateral ventricles (**B**) of wild-type (N=7, A; N=12, B), *Kidins220*^*f/f*^ (N=6, A; N=12, B), and *Kidins220*^*Gfap*Δ/Δ^ (N=10, A; N=15, B), mice. **C**, Cumulative number of cells obtained over 3 consecutive passages of SEZ neurospheres (N=7 independent cultures *per* genotype) and **D**, mean neurosphere diameter of SEZ neurospheres of wild-type (N=9), *Kidins220*^*f/f*^ (N=7) and *Kidins220*^*Gfap*Δ/Δ^ (N=8) mice. **E**, Representative confocal micrographs of wild-type and *Kidins220*^*Gfap*Δ/Δ^ neurospheres stained for the cell cycle marker Ki67 (red) and DAPI (blue) to counterstain the nuclei. Scale bar: 10 μm. **F**, Percentage of Ki67^+^ cells in neurospheres of the three genotypes (wild-type, N=3; *Kidins220*^*f/f*^, N=2; *Kidins220*^*Gfap*Δ/Δ^, N=3). **G**, Representative FACS dot-plots of NSC double-stained for Annexin V/7-AAD and **H**, quantitative analysis of the percentage of live cells (Annexin V^-^/7AAD^-^) and **I**, of early-stage apoptotic cells (Annexin V^+^/7AAD^-^) in wild-type (N=14), *Kidins220*^*f/f*^ (N=6) and *Kidins220*^*Gfap*Δ/Δ^ (N=11) NSCs. Data represent mean ± s.e.m. Each data point represents values from a neurosphere culture established from an individual mouse. Ns, not significant, \**P* < 0.05, ** *P* < 0.01, *** *P* < 0.001, by one-way ANOVA (A, B, D, F, H, I), and two-way RM ANOVA (C), all followed by Dunnett’s *post-hoc* tests. 7AAD, 7-aminoactinomycin D; FACS, fluorescence-activated cell sorting.

### Kidins220 is necessary to suppress GSK3-mediated NSC death, downstream of growth factor signaling

The interaction of various growth factors with their respective receptor tyrosine kinases activates the PI3K/AKT pro-survival pathway. Activated AKT can then phosphorylate the two GSK3 isoforms on their N-terminus (at Ser21 for GSK3α and Ser9 for GSK3β), inactivating them [37, 38]. Suppression of GSK3 activity by PI3K/AKT signaling is indeed the major pathway through which its pro-apoptotic roles are prevented [39]. Given the survival defect of *Kidins220*^*Gfap*Δ/Δ^ neurospheres, we investigated whether the PI3K/AKT signaling pathway was specifically affected by performing immunoblots for the activating phosphorylation of AKT at serine 473 (AKT^pSer473^) [40] and subsequent phosphorylation of GSK3α/β^pSer21/9^. We found a significant reduction in active AKT as well as in the inhibitory phosphorylation of GSK3 isoforms in *Kidins220*^*Gfap*Δ/Δ^ neurospheres (Fig. 5A-C). Given the severe deficit in hippocampal progenitors, neuroblasts and newborn neurons observed *in vivo*, we hypothesized an over-activation of GSK3 in the hippocampus of Kidins220 genetic models and observed a marked deficiency of inactivating phosphorylations in both *Kidins220*^*f*/f^ and *Kidins220*^*Gfap*Δ/Δ^ hippocampal lysates compared to wild-types (Fig. 5D, E). These data indicate that Kidins220 fine-tunes the PI3K/AKT pro-survival pathway in response to external signals in NSCs.

**Figure 5.**
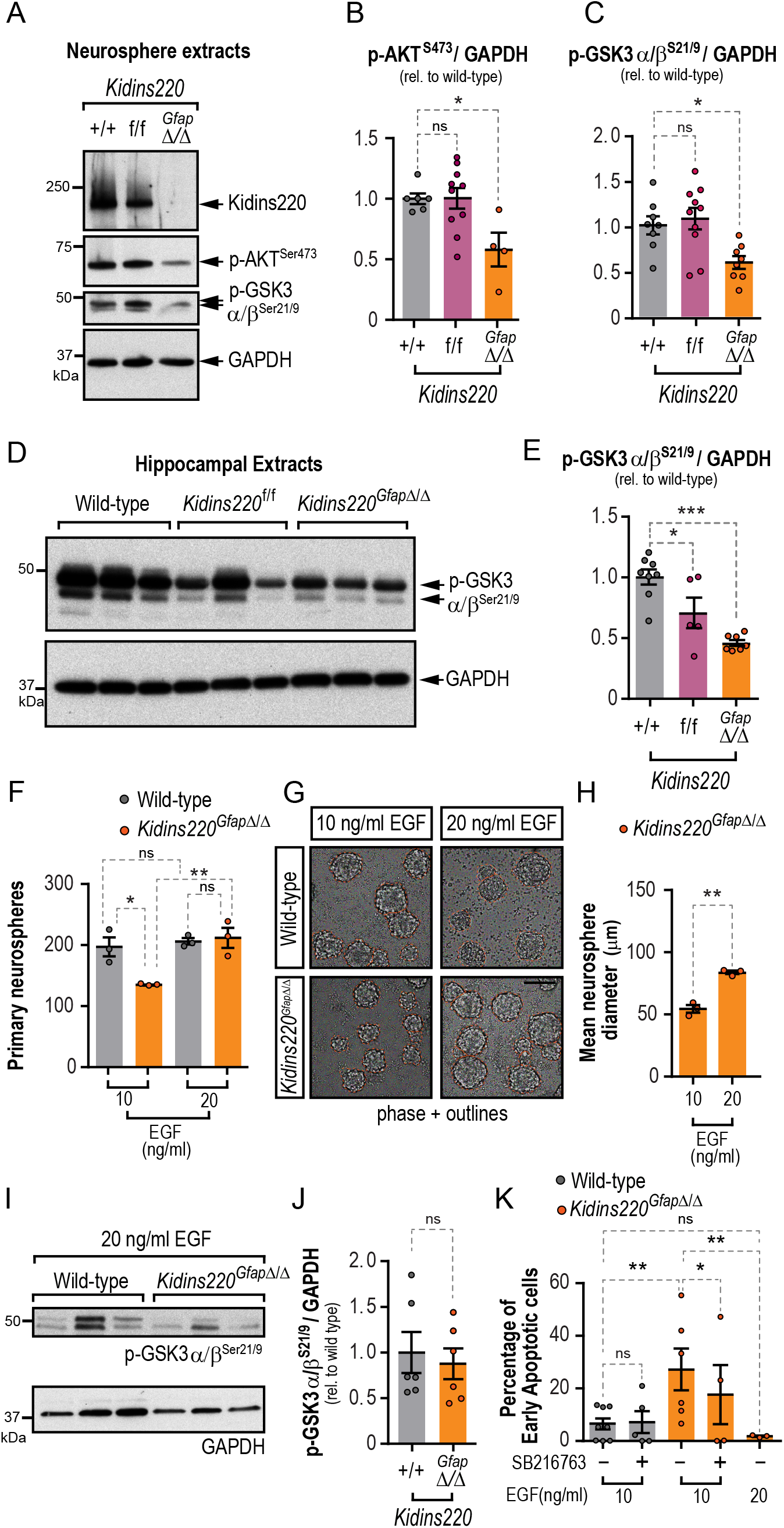
Recuperation of GSK3 inhibition restores survival defect in Kidins220-null neural stem cells. **A**, Representative immunoblots for Kidins220, p-AKT^S473^, p-GSK3α/β^S21/9^ and GAPDH in lysates from *Kidins220*^*Gfap*Δ/Δ^, *Kidins220*^*f/f*^ and wild-type neurospheres. Levels of p-AKT^S473^ (wild-type, N=6; *Kidins220*^*f/f*^, N=10; *Kidins220*^*Gfap*Δ/Δ^, N=3) (**B**) and p-GSK3α/β^S21/9^ (wild-type, N=8; *Kidins220*^*f/f*^, N=10; *Kidins220*^*Gfap*Δ/Δ^, N=8) (**C**) represented in arbitrary units after normalization with GAPDH, and relative to wild-type in extracts from the indicated genotypes. **D**, Representative immunoblots for p-GSK3α/β^S21/9^ and GAPDH in hippocampal lysates from wild-type, *Kidins220*^*f/f*^ and *Kidins220*^*Gfap*Δ/Δ^ mice. **E**, Levels of p-GSK3α/β^S21/9^ represented in arbitrary units after normalization with GAPDH and relative to wild-type in extracts from the indicated genotypes (wild-type, N=8; *Kidins220*^*f/f*^, N=5; *Kidins220*^*Gfap*Δ/Δ^, N=7). **F**, Numbers of primary neurospheres obtained from wild-type and *Kidins220*^*Gfap*Δ/Δ^ mice under low (10 ng/ml) or high (20 ng/ml) EGF mitogenic stimulation (N=3, for each condition). **G**, Representative phase contrast micrographs of wild-type and *Kidins220*^*Gfap*Δ/Δ^ primary neurospheres grown in culture medium with EGF 10 or 20 ng/ml. Neurosphere borders are outlined in orange for clarity. Scale bar 100 μm. **H**, Quantification of the diameters of *Kidins220*^*Gfap*Δ/Δ^ neurospheres grown under low and high EGF (N=3, for each condition). **I**, Representative immunoblots for p-GSK3α/β^S21/9^ and GAPDH in lysates from *Kidins220*^*Gfap*Δ/Δ^ and wild-type primary neurospheres grown in 20 ng/ml EGF. **J**, Levels of p-GSK3α/β^S21/9^ represented in arbitrary units after normalization with GAPDH and relative to wild-type in extracts from the indicated genotypes (N=6, for each condition). **K**, Quantification of the percentage of apoptotic cells in 7-AAD and Annexin V staining detected by flow-cytometry. Wild-type and *Kidins220*^*Gfap*Δ/Δ^ neurospheres were grown in 10 or 20 ng/ml EGF and treated with 100 nM SB216763 GSK3 inhibitor or vehicle (N=8, 5: wild-type, 10 ng/ml EGF, vehicle or SB216763, respectively; N=6, 4: *Kidins220*^*Gfap*Δ/Δ^, 10 ng/ml EGF, vehicle or SB216763, respectively; N=3: *Kidins220*^*Gfap*Δ/Δ^, 20 ng/ml EGF). Data represent mean ± s.e.m. Each data point represents values from a neurosphere culture established from an individual mouse (B, C), or brain extracts from an individual mouse (E). Ns, not significant, \**P* < 0.05, ** *P* < 0.01, *** *P* < 0.0001, by one-way ANOVA followed by Dunnett’s (B, C, E) and Bonferroni’s (F) *post-hoc* tests, two-tailed paired (H, K) and unpaired *t*-tests (J, K).

Next, we hypothesized that increasing the concentration of EGF, the main mitogen in neurosphere cultures, could rescue the cell survival defect of *Kidins220*^*Gfap*Δ/Δ^ NSCs. *Kidins220*^*Gfap*Δ/Δ^ SEZ tissue seeded in medium with a higher EGF concentration (20 ng/ml) yielded ∼30% more primary neurospheres than when seeded in low EGF (10 ng/ml), rising to wild-type numbers, which were unaffected by increasing EGF (Fig. 5F). The higher dose of EGF augmented the diameter of *Kidins220*^*Gfap*Δ/Δ^ neurospheres by ∼40% (Fig. 5G, H). To test if enhancing signaling through EGF restored GSK3 inhibitory phosphorylation to wild-type levels, we performed immunoblots in extracts from neurospheres grown in high EGF and found no differences between genotypes (Fig. 5I, J). Next, we analyzed whether increasing EGF concentration could rescue cell viability. Culturing *Kidins220*^*Gfap*Δ/Δ^ cells in 20 ng/ml EGF rescued the enhanced apoptosis found when cultured in 10 ng/ml EGF, decreasing cell death (Fig. 5K). Furthermore, treatment of wild-type and *Kidins220*^*Gfap*Δ/Δ^ cells with the GSK3 inhibitor SB216763 could still circumvent cell apoptosis in *Kidins220*^*Gfap*Δ/Δ^ cells, although to a lesser extent than a high dose of EGF (Fig. 5K).

## DISCUSSION

In this work we have identified Kidins220 as a critical mediator of hippocampal adult neurogenesis, and a novel intrinsic regulator of NSCs and progenitors, that plays a role tuning PI3K/AKT pro-survival signaling in response to mitogenic stimulation. Mechanisms of survival in adult NSC populations are barely known, even less those that may be related to mitogens that stimulate their division, as survival is a prerequisite for proliferation. Our data indicate that the level of Kidins220 sets the threshold of mitogen-induced activation that sustains adult NSC survival.

*Kidins220* is an essential gene and its global deletion causes extensive cell death in the developing nervous system, accompanied by a growth defect, with embryos displaying smaller bodies and brain size [14, 15]. We show that Kidins220 is abundantly expressed in NSCs from the SEZ and SGZ and that hippocampal neurogenesis is drastically dependent on Kidins220 levels. Neurogenesis in the DG starts during embryonic development and continues through adulthood with the addition of newborn granule neurons [41]. Since Kidins220 deletion using a *Nes*-Cre driver does not cause increased cell death in the embryonic hippocampus at E18.5, when cells accumulate in the DG [14, 15], the effect of Kidins220 deficiency on hippocampal development appears to be mainly postnatal. A reduction in Kidins220 markedly decreases type 2 cycling progenitors and the concomitant emergence of DCX+ neuroblasts and adult-born neurons, indicating that survival might be compromised when Kidins220 levels drop in the DG. Accordingly, Kidins220 deletion in immature hippocampal neurons leads to multiple axons and dendritic aberrations [42], whereas its silencing in mature cortical cultured neurons increases their death and vulnerability to excitotoxicity [16, 18]. Notably, the defects we identify herein correlate with impaired hippocampal-dependent spatial working memory, a trait strongly associated with depletion of adult hippocampal neurogenesis [43].

The sharp decline in hippocampal neurogenesis with advancing age is associated to age-related cognitive deficits due to its function in learning and memory. As in Kidins220-deficient mice, neurogenesis at the DG of aged mice is substantially reduced relative to the SEZ [44], consistent with a potential premature aging of neurogenic niches in these models. Importantly, administration of growth factors restored aging-associated neurogenic defects in both niches [44]. Similarly, our data show that *Kidins220*^*GfapΔ/Δ*^ SEZ-derived neurospheres are rescued by extrinsic cues, as their number, size, and pro-survival pathways are restored to wild-type values by increasing the concentration of EGF.

The extreme sensitivity of DG neurogenesis to reduced Kidins220 levels is in contrast with our findings from the SEZ, which seems refractory to Kidins220 deficiency. This fact may indicate that, in addition to the cell autonomous effect described, the hippocampal niche may be altered in Kidins220-deficient mice and does not provide the adequate cues, a possibility that we have not explored herein. In this regard, Kidins220 expression in astrocytes is required for the control of Ca^(2+)^ dynamics, and thus, it mediates the establishment of proper neuronal connectivity [45]. Our data support the notion that Kidins220 is crucial for cell survival of neurons and precursors, without majorly affecting their proliferation. Neurotrophin signaling may be key in Kidins220-mediated control of DG ontogenesis and adult neurogenesis, since Kidins220 has been shown to interact with all Trk-receptors and with p75NTR [46, 47], and reduced neuronal numbers may be due to decreased trophic support in Kidins220-deficient mice by diminished response to neurotrophins.

Kidins220-deficient mice present hydrocephalus [23], and because previous reports have linked reduced SEZ NSCs numbers to this condition *in vivo* [48, 49], we have only investigated mice with moderate enlargement of the ventricles (see Fig. S3) in this study. Our data suggest that the milder presentations of ventriculomegaly do not affect SEZ NSCs and progeny, at least in our Kidins220-deficient mice models and at the age studied. Our data call for further investigations on the SEZ in Kidins220-deficient mice afflicted with severe hydrocephalus, as future follow-up for this work.

Keeping Kidins220 levels over a certain threshold maintains PI3K/AKT-mediated GSK3 inhibition in NSCs and hippocampal extracts in both Kidins220-deficient mice strains, and increased dosage of EGF rescues cell survival in NSCs restoring GSK3 inhibition. EGFR does not co-immunoprecipitate with Kidins220 [50], so the possibility of Kidins220 regulating clustering of this receptor in NSCs is unlikely. However, Kidins220 may modulate a downstream EGFR effector. Remarkably, Kidins220 provides a molecular docking site for the CrkL adaptor protein to sustain mitogen-activated protein kinases<u>/extracellular signal-regulated kinases</u> signaling in neurons in response to nerve growth factor [51]. A parallel mechanism might play a role in NSCs, as CrkL directly binds to and regulates PI3K/AKT [52-54]. Since silencing Kidins220 does not alter AKT activation in cultured neurons [47], Kidins220 may be crucial for PI3K/AKT activation in NSCs and neuronal progenitors, in contrast to terminally differentiated neurons.

A growing body of evidence points to a direct role of KIDINS220 in cancer [55-57]. In prostate cancer, KIDINS220 activates the PI3K/AKT pathway [58], and in pediatric high grade glioma copy number breakpoint within *KIDINS220* gene have been identified [57]. NSCs of the SEZ are considered glioma cells of origin [59, 60], and molecular alterations of the EGFR/PI3K/Akt/mTOR axis are hallmarks of this cancer type, which has led to the design of clinical trials devised to target this pathway [61]. Whether KIDINS220 could be part of the stimulus-response threshold activating this pro-survival pathway downstream of tyrosine kinase receptors in glioma stem cells and possible therapeutic implications, deserves further investigation.

Maintaining the proper levels of GSK3 activity is crucial for neural progenitor maintenance, proliferation, survival, and differentiation [62, 63]. While GSK3 inhibition promotes adult hippocampal neurogenesis [64, 65], increased GSK3 activity reduced DG volume and causes defects in neuroblast generation, features like those reported herein [66]. Consistently, failure to properly regulate GSK3 activity plays a role in Alzheimer’s disease and schizophrenia [67-69], diseases associated with Kidins220 alterations, amongst others [20, 22-25, 70]. Exciting new studies support the idea that neurogenic potential preservation may be crucial to restrain cognitive decline associated with neurodegenerative conditions and non-physiological aging in humans [71, 72]. The data presented herein point to the possibility that alterations in KIDINS220 could render the DG neurogenic niche more sensitive to cognitive decline, and memory and learning disabilities, traits associated with these diseases. In summary, our data pinpoints Kidins220 as a multifaceted molecule with relevant functions in NSCs and adult neurogenesis. Dissecting the molecular basis of NSCs activation to generate new progeny grants potential for conceiving novel therapeutic strategies based in stem cells.

## MATERIALS AND METHODS

### Mice

*Kidins220*^*f*/f^ mice have been described previously (21, 29). B6.Cg-Tg(Gfap-cre)73.12Mvs/J (herein referred to as Gfap-Cre) were purchased from The Jackson Laboratory, (Bar Harbor, Maine, USA; JAX stock #012886) [32]. Housing of mice and all experiments were conformed to the appropriate national legislations (RD 53/2013) and the guidelines of the European Commission for the accommodation and care of laboratory animals (revised in Appendix A of the Council of Europe Convention ETS123), following protocols approved by the corresponding local ethics committees. Male and female 2-month-old *Kidins220*^*f*/f^, *Kidins220*^*Gfap*Δ/Δ^ or wild-type littermates were employed. Genotyping and recombination were monitored in genomic DNA samples by PCR using specific pairs of primers (Table S1). For long term label retention experiments, mice were injected and brains collected and processed, or fresh tissue was dissected to obtain whole mounts of the SEZ as described elsewhere [30, 31].

### Immunohistochemistry of mouse brain samples

Immunostainings were performed in free-floating sections, incubated for 48 h with the appropriate primary antibodies (Table S2). For the specific detection of the synthetic nucleosides, a 20 min 2 N HCl denaturalization followed by neutralization in borate buffer was performed before the addition of the antibodies. Immunofluorescent detections were performed with Alexa Fluor (Invitrogen) conjugated secondary antibodies. DAPI (1 μg/ml) was used for counterstaining. Images were acquired under the same settings for each experiment (Fluoview FV10i Olympus, Shinjuku, Tokyo, Japan, and Zeiss LSM 710, Zeiss, Germany confocal laser scanning microscopes, equipped with 405, 458, 488 and 633 nm lasers. Images were acquired and processed using FV10-ASW 2.1 viewer (Olympus) and ImageJ/FIJI (version 1.33; National Institutes of Health, Bethesda, MD) software.

### Cell Counting and dentate gyrus volume

Numbers of NSC at the ventricle surface, CldU+ cells in the SEZ, the OB and the corpus callosum were obtained as described previously [31, 73]. The total volume of the DG and the number of CldU^+^-GFAP^+^ NSC, precursor cells (IdU^+^-Sox2^+^), immature neurons (Dcx^+^), and mature neurons (CldU^+^-NeuN^+^) were calculated using the physical dissector method adapted for confocal microscopy as described elsewhere [73]. Briefly, DG area was measured in each slice using ImageJ software in one confocal series composed of every fifth section of the whole DG. The DG volume was calculated as the area obtained × thickness of the slice (50 μm), and the total volume of the DG was calculated as the summatory of the volume ×5. Numbers of positive cells for each marker were scored in every plane of one confocal series composed of every sixth section of the whole DG. To obtain total cell numbers, data were multiplied ×6.

### Behavioral tests

Habituation was performed by handling 5 min every other day on the previous week to the test. For the T maze test, mice were food restricted for 48h before the beginning of the test which consisted of an acquisition phase and a reversal phase. Only one of the two short arms was rewarded with food. For each trial, the mouse was placed in the ‘start’ section of the long arm and given a maximum of three minutes to complete the test. In the second phase of the experiment, reversal learning was tested and the contingency was reversed, i.e., the other short arm was rewarded. Acquisition and reversal phases lasted 5 days (6 daily trials, with a two-day break between both phases). We scored the number of visits to the non-rewarded arm (wrong visits), the time to reach the reward and start eating (latency), and accuracy (1 point was given for entering the rewarded arm and eating the pellet within 30 seconds; any other option was scored with 0 points).

### Magnetic resonance imaging

Magnetic resonance imaging (MRI) was performed as described (21). Briefly, MRI studies were performed in a Bruker Biospect 7.0-T horizontal-bore system (Bruker Medical Gmbh, Ettlingen, Germany), equipped with a 1H selective birdcage resonator of 23 mm and a Bruker gradient insert with 90 mm of diameter (maximum intensity 36 G/cm). Data were acquired using a Hewlett-Packard console running Paravision software (Bruker Medical Gmbh) operating on a Linux platform.

### Cell culture, immunofluorescence and cell viability analysis by flow cytometry

Neurosphere cultures were established and maintained as described [34, 35] or were cultured in adhesive conditions as described previously [36] using Geltrex (Invitrogen, Waltham, MA, USA,) as coating substrate, with murine natural EGF (10 or 20 ng/ml, Thermofisher Scientific, Waltham, MA, USA) and human recombinant FGF-2 (10 ng/ml; Merck Millipore, Burlington, MA, USA) as mitogens. Primary neurosphere numbers and diameters were assessed by manual counting and photographed in an inverted microscope (Axiovert200, Zeiss ORCA-Flash4.0 LT sCMOS, Hamamatsu Photonics, Hamamatsu, Japan), and diameters measured using ImageJ/FIJI (NIH). For immunofluorescence, neurospheres were allowed to attach to Geltrex for 10 min, fixed with PFA 4%, and antibody staining was performed as previously described [74]. Culture of primary cortical astrocytes was performed as described [23]. For flow cytometry, and rescue experiments, cells were cultured with 10 or 20 ng/ml EGF for three passages before cells were seeded on Geltrex-coated plates with the appropriate dose of EGF and treated with DMSO or 100 nM SB216763 (Selleckchem, Houston, TX, USA) for 3 days. Cells were harvested by centrifugation, disaggregated, and resuspended in Annexin V Binding Buffer, PE Annexin V and 7-AAD (BD Iberia, Madrid, Spain) and analyzed in a FACSCanto-A flow cytometer (BD) using FACSDiva 6.1.3 software (BD). Data were analysed with FlowJo software (Tree Star).

### Preparation of protein extracts and immunoblot analyses

Cell extracts were obtained by lysis in SDS-buffer (25 mM Tris-HCl, pH 7.4, 1 mM EDTA, 1% SDS) or in RIPA buffer (25 mM Tris-HCl, pH 7.6, 1% Triton X-100, 0.5% sodium deoxycholate, 0.1% SDS, 150 mM NaCl) with protease and phosphatase inhibitors for 30 min at 4 °C. Lysates were centrifuged for 30 min at 14 000 rpm at 4°C, and supernatant was considered the total lysate or total lysate soluble fraction. Lateral ventricles including the SEZ, and whole hippocampi were dissected, frozen in dry ice and homogenized in RIPA buffer as above using potter and Polytron homogenizers. Protein concentration was determined with BCA Protein Assay kit (Pierce Biotechnology Inc., Waltham, MA, USA); 30-100 μg of protein were resolved by SDS-PAGE and transferred to nitrocellulose membranes (Protran, Sigma-Aldrich). Membranes were blocked in PBS or TBS containing 3% BSA or 5% skimmed milk, and 0.025% Tween-20 and incubated with primary antibodies (Table 2), followed by appropriate peroxidase-conjugated anti-mouse or anti-rabbit IgGs (Dako). Signal was obtained by enhanced chemiluminescence (Western Lightning. Perkin Elmer, <u>Waltham, MA,</u> USA). Auto-radiographic films were scanned, and the bands were analyzed by densitometry using Image-J software (NIH). Immunoblot images have been cropped for presentation.

### Statistical analyses

Analyses of significant differences between means were performed with Prism 9 software (GraphPad, San Diego, CA) as specified in each figure legend. Results are shown as mean ± s.e.m. and the number of experiments (N) carried out with independent subjects (primary cultures or mice) is shown in each figure as dots and is specified in the figure legends. Data transformations using square root, or arcsin were performed to meet test assumptions (normality and/or homoscedasticity) when needed. In all cases, *P* < 0.05 denoted statistical significance. Appropriate *post-hoc* tests were applied when necessary. No statistical method was used to predetermine sample size. Sample sizes were based on previous published experiments

## Supporting information

del Puerto et al Supplemental Information

## ACKNOWLEDGMENTS

We are grateful to Professor R. Fernández-Chacón (IBIS, Sevilla), and J.A. Morales-García and A. Perez-Castillo (IIBM, CSIC-UAM) for their help with the hippocampal cultures, and A. Pérez-Villalba (UCV, Valencia) for suggestions with behavior analyses. We acknowledge all members of our laboratories for constructive suggestions and the excellent technical support by S. Benito, B. Ortigosa, M. Prudencio Sánchez-Carralero and A. González-Martín. Our special thanks to L. Sánchez-Ruiloba for her assistance in confocal images analysis, and to the specialized support of core facilities personnel at the IIBM (CSIC-UAM): animal care, SIERMAC, confocal microscopy; and at the CBMSO (CSIC-UAM): confocal microscopy and flow cytometry.

## AUTHOR CONTRIBUTIONS

AdP, BM-P, ALB-M, CL-F, JP-U, BA-S and EP performed research. FC and GS generated Kidins220 floxed mice. MR-C and IF provided reagents and tools. All authors contributed to data interpretation and analysis and to the writing of the manuscript. TI and EP performed study concept and design, secured funding, supervised the work and conceived and wrote the manuscript.

## ETHICS

Ethics statement regarding animal procedures is included in Materials and Methods section.

## DATA AVAILABILITY

All data are available under reasonable request.

