## Supplementary material for "Kidins220 sets the threshold for survival of neural stem cells and progenitors to sustain adult neurogenesis": del Puerto et al Supplemental Information

**This PDF file includes:**

Figures S1 to S4
Tables S1 to S3

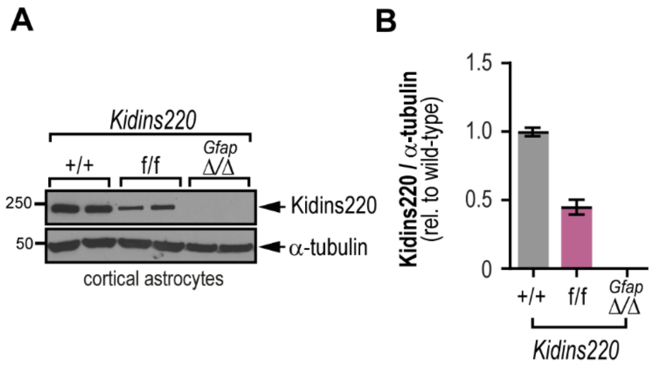


**Fig. S1. Confirmation of the full excision of the floxed allele in astrocytes from *Kidins220*$^{Gfap\Delta/\Delta}$ mice. A,** Immunoblot for Kidins220 and α-tubulin (loading control) in primary cortical astrocytes from the denoted genotypes. **B**, Kidins220 levels in arbitrary units after normalization with α-tubulin in wild-type and Kidins220 genetically modified mice. Data represent mean ± s.e.m. (N=2, for each genotype).

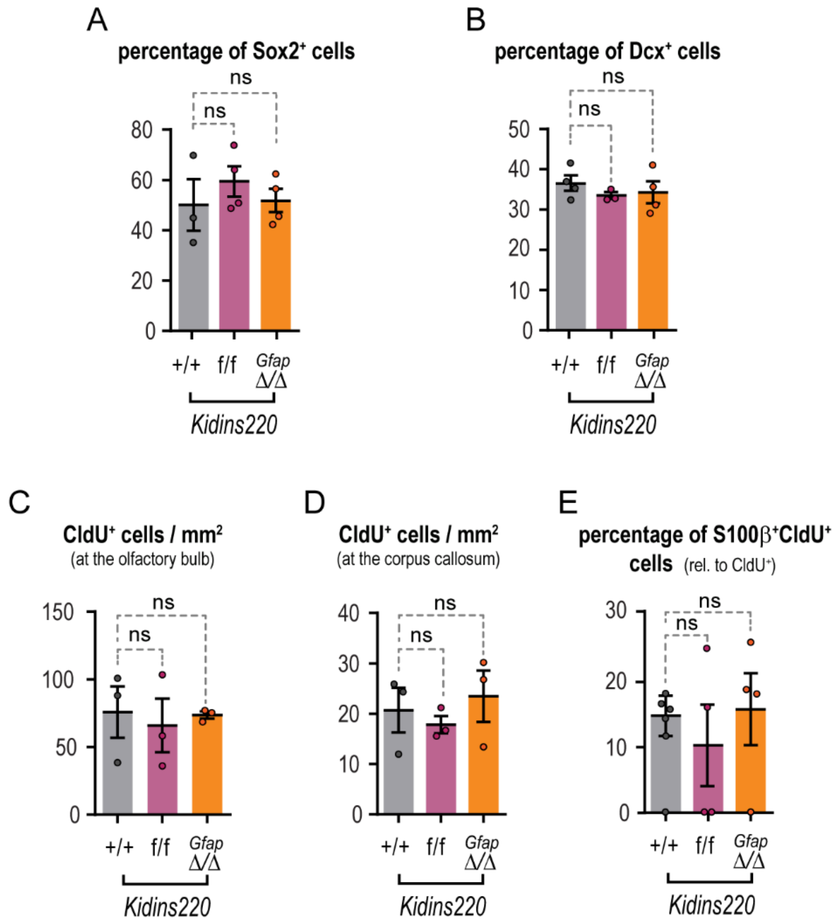


**Fig. S2. Adult cytogenesis from the 2 month-old SEZ and *corpus callosum* is Kidins220 independent.** Percentage of Sox2$^+$ (**A**) and Doublecortin (DCX)$^+$ (**B**) in the subependymal zone of 2 month-old wild-type and Kidins220 genetically modified mice. Sox2$^+$-ependymal cells were excluded from the analysis. Mice were injected with CldU as described in methods section and sacrificed 28 days later. CldU$^+$ cells in the glomerular layer of the OBs (**C**) and in the *corpus callosum* (**D**) were scored. **E**, CldU$^+$ cells, double positive for the astrocytic marker S100-ß were quantified in the SEZ of wild-type and Kidins220 genetic models. Graphs represent mean ± s.e.m. where each data point denotes an individual mouse (N=3-5, for each genotype). Ns, not significant. One-way ANOVA, followed by Dunnett's *post-hoc* test.

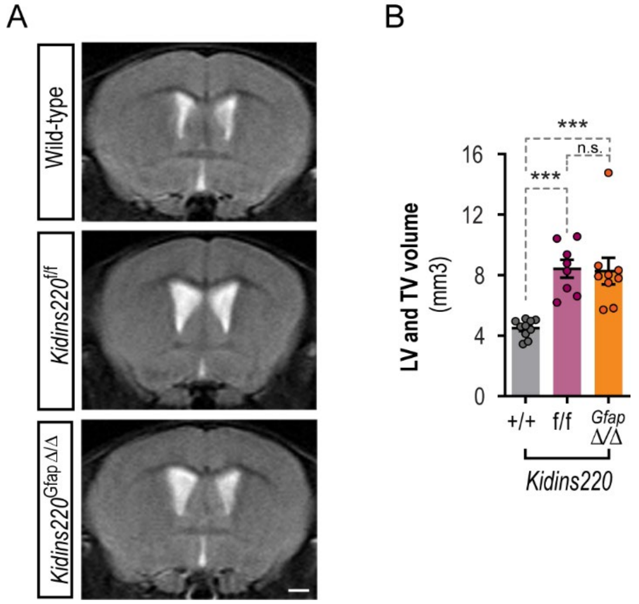


**Fig. S3. *Kidins220$^{Gfap\Delta/\Delta}$* mice present ventriculomegaly. A,** Representative *in vivo* T2-weighted (T2-W) MRI coronal images showing lateral and third ventricles (LVs + TV) of 2-month-old wild-type, *Kidins220$^{f/f}$* and *Kidins220$^{Gfap\Delta/\Delta}$* male mice. Scale bar, 1 mm. **B**, Quantification of LV and TV ventricular volume of 10 Wild-type, 8 *Kidins220$^{f/f}$* and 9 *Kidins220$^{Gfap\Delta/\Delta}$* mice. Graph represents mean ± s.e.m. where each dot denotes values from an individual mouse (N=8-10, for each genotype). Ns, not significant, *** P < 0.001, by one-way ANOVA, followed by Tukey's *post-hoc* test.

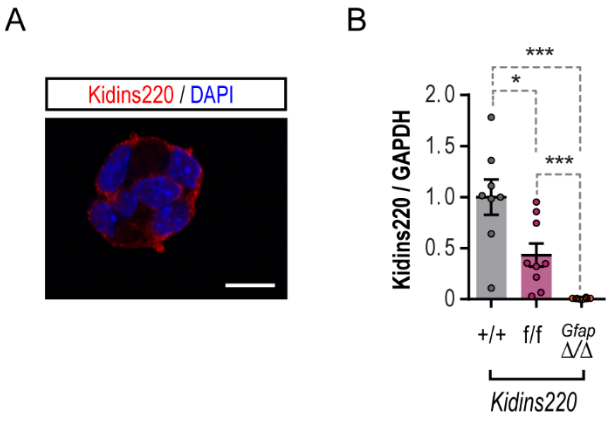


**Fig. S4. Kidins220 expression in neurospheres from wild-type and Kidins220 genetic models. A**, Kidins220 (red) is highly expressed in neurospheres at the cellular membrane. Nuclei are stained with DAPI (blue). Scale bar, 10 μm. **B,** Kidins220 levels represented in arbitrary units after normalization with GAPDH and relative to wild-type in extracts from wild-type, *Kidins220*$^{f/f}$ and *Kidins220*$^{Gfap\Delta/\Delta}$ mice t after immunoblot analysis of lysates from neurospheres obtained from lateral ventricle walls. Data represent mean ± s.e.m. Each data point represents a neurosphere culture established from an independent mouse (N=8-9, for each genotype). **P* < 0.05, *** *P* < 0.0001 by one-way ANOVA followed by Tukey's *post-hoc* test.

**Table S1. Sequences of oligonucleotides used for genotyping**

| *Sequence* | *Use* |
|---|---|
| *5'- GAGCACAGACTTCTCTTATGG -3'* | Forward primer for detection of the *Kidins220*-floxed and recombined alleles |
| *5'- GCGTTTCTAGCATACACATG -3'* | Reverse primer for detection of the *Kidins220* floxed allele |
| *5'-CAGATGGCTGTGAACCACCGTTTAAAC-3'* | Reverse primer for detection of the *Kidins220* recombined allele |
| *5'- GCGGTCTGGCAGTAAAAACTATC -3'* | Forward primer for detection of *Cre* |
| *5'- GTGAAACAGCATTGCTGTCACTT -3'* | Reverse primer for detection of *Cre* |
| *5'- CTAGGCCACAGAATTGAAAGATCT -3'* | Forward primer for detection of internal control DNA |
| *5'- GTAGGTGGAAATTCTAGCATCATCC -3'* | Reverse primer for detection of internal control DNA |

**Table S2. Primary antibodies used in this work**

| *Antigen* | *Working dilution & use* | *Origin* |
|---|---|---|
| *Kidins220* | 1:500 IF/IHC | (15) from T. Iglesias lab. |
| *Kidins220* | 1:250 IF/IHC | (10) from T. Iglesias lab. |
| *Kidins220* | 1:1000 IB | (10) from T. Iglesias lab. |
| *Sox2* | 1:300 IHC | Abcam ab97959 |
| *Sox2* | 1:250 IHC | R&D Systems AF2018 |
| *Ki67* | 1:200 IF | Abcam ab15580 |
| *IdU* | 1:500 IHC | BD #347580 |
| *CldU* | 1:500 IHC | Abcam ab6326 |
| *GFAP* | 1:500 IHC | Chemicon International ab5541 |
| *γ-Tubulin* | 1:500 IHC | Sigma-Aldrich T6557 |
| *γ-Tubulin* | 1:250 IHC | Abcam ab11317 |
| *S100-β* | 1:100 IHC | Abcam ab52642 |
| *GAPDH* | 1:1000 IB | Merck Millipore MAB374 |
| *α-Tubulin* | 1:10 000 IB | Sigma-Aldrich T9026 |
| *β-Actin* | 1:40 000 IB | Sigma-Aldrich A2228 |
| *p-GSK3α/β*$^{S21/9}$ | 1.1000 IB | Cell Signaling Technology #9331 |
| *p-AKT*$^{S473}$ | 1:1000 IB | Cell Signaling Technology #4060 |
| *AKT* | 1:1000 IB | Cell Signaling Technology #2966S |
| *N-Cadherin* | 1:100 IHC | BD Transduction Laboratories AB_398236 |
| *β-Catenin* | 1:100 IHC | Cell Signaling Technology #9562 |

IF, immunofluorescence; IHC, immunohistochemistry; IB, immunoblot

**Table S3. Two-way RM ANOVA related to Fig. 3A**

| | | Interaction | Genotype | Session |
|---|---|---|---|---|
| **Acquisition Phase** | **Latency** | F (8, 104) = 0.9036<br>P=0.5165 ns | F (3.269, 85.00) = 80.70<br>P<0.0001 *** | F (2, 26) = 1.002<br>P=0.3807 ns |
| | **Accuracy** | F (8, 104) = 1.290<br>P=0.2566 ns | F (3.385, 88.01) = 41.61<br>P<0.0001 *** | F (2, 26) = 0.2263<br>P=0.7990 ns |
| | **Wrong Visits** | F (8, 104) = 1.160<br>P=0.3305 ns | F (3.055, 79.43) = 39.12<br>P<0.0001 *** | F (2, 26) = 0.5582<br>P=0.5790 ns |
| **Reversal phase** | **Latency** | F (8, 104) = 0.8745<br>P=0.5405 ns | F (2.178, 56.63) = 5.403<br>P=0.0058 ** | F (2, 26) = 10.46<br>P=0.0005 *** |
| | **Accuracy** | F (8, 104) = 0.8048<br>P=0.5997 ns | F (3.223, 83.79) = 9.367<br>P<0.0001 *** | F (2, 26) = 16.66<br>P<0.0001 *** |
| | **Wrong Visits** | F (8, 104) = 1.006<br>P=0.4360 ns | F (2.317, 60.24) = 5.948<br>P=0.0029 ** | F (2, 26) = 5.953<br>P=0.0074 ** |
